# Changes in taste palatability across the estrous cycle are modulated by hypothalamic estradiol signaling

**DOI:** 10.1101/2024.04.01.587593

**Authors:** Bradly T. Stone, Oran M. Rahamim, Donald B. Katz, Jian-You Lin

**Affiliations:** Neuroscience Program, Brandeis University, Waltham, MA, USA; Department of Psychology, Brandeis University, Waltham, MA, USA; Volen National Center for Complex Systems, Brandeis University, Waltham, MA, USA

## Abstract

Food intake varies across the stages of a rat’s estrous cycle. It is reasonable to hypothesize that this cyclic fluctuation in consumption reflects an impact of hormones on taste palatability/preference, but evidence for this hypothesis has been mixed, and critical within-subject experiments in which rats sample multiple tastes during each of the four main estrous phases (metestrus, diestrus, proestrus, and estrus) have been scarce. Here, we assayed licking for pleasant (sucrose, NaCl, saccharin) and aversive (quinine-HCl, citric acid) tastes each day for 5-10 days while tracking rats’ estrous cycles through vaginal cytology. Initial analyses confirmed the previously-described increased consumption of pleasant stimuli 24-48 hours following the time of high estradiol. A closer look, however, revealed this effect to reflect a general magnification of palatability—higher than normal preferences for pleasant tastes and lower than normal preferences for aversive tastes—during metestrus. We hypothesized that this phenomenon might be related to estradiol processing in the lateral hypothalamus (LH), and tested this hypothesis by inhibiting LH estrogen receptor activity with ICI_182,780_ during tasting. Control infusions replicated the metestrus magnification of palatability pattern; ICI infusions blocked this effect as predicted, but failed to render preferences “cycle free,” instead delaying the palatability magnification until diestrus. Clearly, estrous phase mediates details of taste palatability in a manner involving hypothalamic actions of estradiol; further work will be needed to explain the lack of a flat response across the cycle with hypothalamic estradiol binding inhibited, a result which perhaps suggests dynamic interplay between brain regions or hormones.

**Significance Statement:** Consummatory behaviors are impacted by many variables, including naturally circulating hormones. While it is clear that consumption is particularly high during the stages following the high-estradiol stage of the rodent’s estrous (and human menstrual) cycle, it is as of yet unclear whether this phenomenon reflects cycle stage-specific palatability (i.e., whether pleasant tastes are particularly delicious, and aversive tastes particularly disgusting, at particular phases). Here we show that palatability is indeed modulated by estrous phase, and that this effect is governed, at least in part, by the action of estradiol within the lateral hypothalamus. These findings shed light on the mechanisms underlying the adverse impact on human welfare due to irregularities observed across the otherwise cyclic menstrual process.

## Introduction

Hormones play critical roles in determining an animal’s behavioral responses to internal and external stimuli. Notable among the behaviors impacted by hormones are those associated with feeding: fluctuations of reproductive hormone levels impact what, and how much, food and drink a cycling animal consumes (Asarian & Geary, 2013; Duval et al., 2014; Eckel, 2004; Martini et al., 1994; Rivera & Stincic, 2018). In humans, for instance, consumption of energy- and nutrient-rich diets increases during the midluteal phase of the menstrual cycle (relative to the midfollicular phase; Martini et al., 1994). Similar phenomena have been observed in relation to the (analogous) rat 4-day estrous cycle, in that rats consume more following estradiol peaks (i.e., during metestrus and diestrus) than they do during phases leading up to these peaks (proestrus/estrus; Fantino & Brinnel, 1986; Ter Haar, 1972).

It is reasonable to hypothesize that shifts in taste palatability—how delicious or disgusting the animal finds a substance in its mouth—might underly such hormone-related fluctuations in food intake. But while a subset of studies has supported this view, the evidence has been less than conclusive: a few research groups have suggested, on the basis of real-time tests of licking or orofacial responsiveness, that phase specificity of sucrose and NaCl consumption is indeed palatability-related (Atchley et al., 2005; Curtis et al., 2004; Pereira Jr et al., 2019; Rivera et al., 2007); others suggest, however, that the time course of this preference development (i.e., the fact that it does not appear in the first minute of testing) implicates learning rather than basic perceptual effects (Hrupka et al., 1997), or that the phenomenon reflects a post-ingestive (and therefore post-taste) mechanism (Eckel, 2004; Geary et al., 1995).

One possible reason that the above-described experiments are inconclusive lies in their reliance on consumption data for single (usually pleasant) tastes. Palatability is an intrinsically relative construct, and therefore changes in consumption of single tastes are difficult to interpret in the absence of comparisons to consumption of tastes with different palatabilities. Only a few studies have performed such comparisons: Clarke and Ossenkopp (1998) examined both positive and negative taste reactivity in female rats, and found only mixed (depending on the specific behavior analyzed) evidence for palatability processing differences between high and low estradiol halves of the 4-day cycle; Parker et al. (2002), meanwhile, reported a mild increase in consumption of sugary cereal during broadly-defined diestrus (a period that may well have included metestrus, which is relatively brief; see below and Butcher et al., 1974; Cora et al., 2015), without a comparable increase in consumption of less pleasant (non-sweet) chow. Overall, while the evidence is suggestive, more work testing this hypothesis is clearly needed (a point recently made by Rivera & Stincic, 2018).

Of course, dissecting the ways in which palatability-related behavior changes across estrous phases is a non-trivial problem, because rodents cycle fast—each phase lasts approximately 1 day. One way to deal with this technical difficulty would be to perform tests on ovariectomized rats, such that circulating estradiol levels can be determined by experimenter injection (Taschetto et al., 2018; Yoest et al., 2019; Yokota-Nakagi et al., 2022). We have chosen not to use this technique here, because while it allows nominal control of estradiol levels (in isolation of other sex-specific hormones), it is difficult to be sure that one is administering the hormone of interest at naturalistic concentrations (studies tend to involve estradiol injections in the vicinity of 5µg/0.1mL, versus natural fluctuations between 2-50pg/mL across the cycle; see for instance Yoest et al., 2019). By first performing quick assessments of behavior in more naturalistic conditions, an approach that has long been the choice of researchers interested in how behavior fluctuates across estrous phases (e.g., Atchley et al., 2005; Parker et al., 2002), we can make fewer potentially problematic assumptions about how hormone infusions spread through the system, about how hormones do (or don’t) interact, and about across-cycle subtleties of hormone dynamics. By allowing hormones to vary in their normal range, it is easier to distinguish an important hormone from an all-important hormone, ultimately ensuring that the results of later ovariectomy experiments will be maximally interpretable.

We therefore employed the Brief Access Task (BAT; Smith, 2001), with which preferences toward a battery of pleasant and aversive tastes can be quantified and compared within single test sessions (and cycle phases). This speed of assessment that we were not limited to making “relatively high vs. relatively low consumption” comparisons of the two halves of the cycle— instead we could identify individual estrus phases that were uniquely different from the other three, and thereby observe a wholistic magnification of the range of palatabilities (that is, rats simultaneously showed higher than normal preference to pleasant tastes and lower than normal preference to aversive tastes) centered specifically on metestrus.

We hypothesized that this phase-specific palatability modulation might reflect a centrally-mediated action of estradiol. Among the many brain regions that are rich in estrogen receptors (ERs; Santollo & Daniels, 2015a, 2015b), the lateral hypothalamus (LH) was selected as the target of interest, because LH is deeply implicated in the modulation of consumption (Fujisawa et al., 2001; Touzani & Velley, 1990) and palatability processing (Li et al., 2013), and because LH is directly connected to both taste and reward circuits (Berthoud & Münzberg, 2011; Fadel & Deutch, 2002) that are also critically involved in the modulation of food consumption (Fu et al., 2019; Petrovich et al., 2005; Tyree & de Lecea, 2017). To test this hypothesis, we inhibited hypothalamic ERs while repeating the experiment, a manipulation that did, in fact, block the metestrus effect. Rather than rendering preferences ’cycle free,’ however, LH estradiol blockade instead delayed preference magnification to the diestrus phase, thereby revealing complexities of the hormonal control of feeding.

Overall, these results demonstrate that the estrous cycle modulates taste palatability, in part through estradiol activity in LH. Further work will be necessary to disentangle the factors— additional brain areas or hormones, perhaps—involved in the influences of estrus cycle on taste preferences that emerges in the absence of hypothalamic estrogen receptor function.

## Results

### Female rats progress through recognizable estrous phases

We evaluated estrous phase by quantifying distinct, specific ratios of cell types (Cora et al., 2015) present in photomicrographs of plated vaginal smears (**Figure 1A-1D**). This quantification was performed visually by a pair of raters who were blind to both the behavioral data and the other rater’s assessments. The inter-rater congruence for identifying estrous phase averaged 76.4%; agreement between the two raters’ assessments (Cohen’s Kappa κ=0.672) is in the acceptably high range (McHugh, 2012). Data from rats that experienced an irregular cycle, or for which agreement could not be easily reached, were removed from analyses.

**Figure 1.**
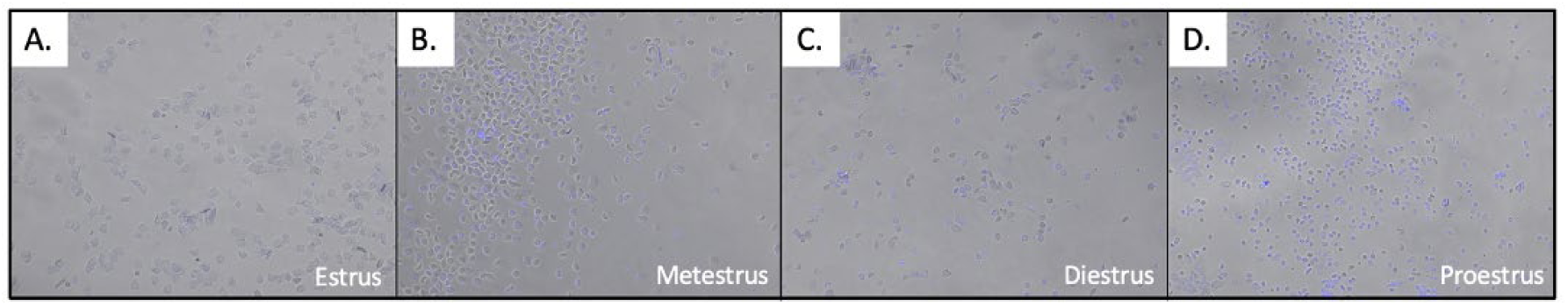
Photomicrographs from representative wet, DAPI-stained vaginal smears obtained from four consecutive test days from a Long Evans rat. Images are in order of estrous cycle with presence of estradiol increasing from left to right: A. Estrus; classic cornified cells, B. Metestrus; presence of a combination of cornified, nucleated, and leukocytes, C. Diestrus; leukocytes appear in combination with nucleated cells, and D. Proestrus; majority of nucleated cells appear in clumps. Note. The inter-rater congruence for identifying estrous phase; Overall Cohen’s *κ* confirmed substantial inter-rater reliability (*κ* = 0.672).

### Relative preference for pleasant tastes depends on estrous phase

Our central question was whether and how the palatabilities of positive and aversive tastants change with estrous phase. Because initial ANOVAs revealed neither a significant effect of Taste nor a Taste x Phase interaction within valence categories (i.e., between pleasant sucrose, saccharin, and NaCl, and between aversive quinine and citric acid, *ps* > 0.05), we increased statistical power by combining these data (**Figure 2**).

**Figure 2.**
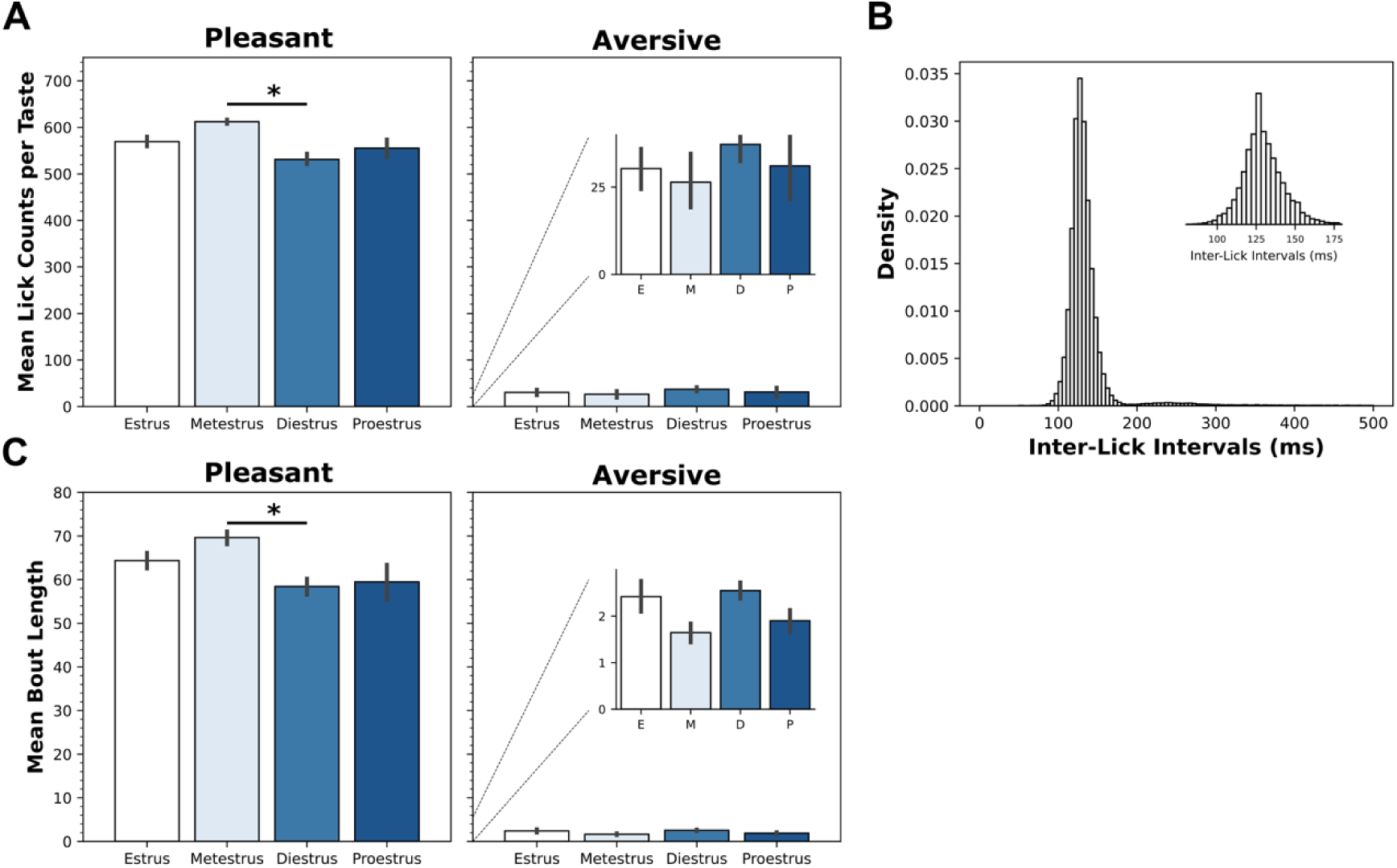
Average intake and lick bout length changes with estrous phase. **A**. Measured in terms of raw lick counts, rats consumed more of the pleasant tastes during metestrus than diestrus; lick counts to aversive tastes did not differ significantly across phases (see text) but “mirrors” those of pleasant tastes (the inset expands the y axis, facilitating comparison of left- and right-panel patterns). **B**. Histogram of inter-lick intervals from a representative rat is normally distributed with a mean of ∼125 ms, demonstrating stereotypical rodent licking behavior (lick rate ∼8 Hz); data from the entire group (N = 10) are shown in inlay. **C**. Rats produced longer bouts in response to pleasant tastes during metestrus than diestrus; as with lick counts, bout lengths to aversive tastes didn’t differ between phases (again, the inset expands the y axis, facilitating comparison of left- and right-panel patterns). * *p* < 0.05. Error bars represent SEMs. E: Estrus; M: Metestrus; D: Diestrus; P: Proestrus.

One-way ANOVAs performed separately on pleasant and aversive stimuli revealed that the consumption of pleasant tastes is impacted by estrous phase (**Figure 2A-left**, *F*(3,146)=4.19, *p* < 0.01). Post hoc comparisons (*Turkey-HSD* tests) clarified that rats consume significantly less pleasant tastant during diestrus than metestrus (*p* < 0.05); no other between-phase differences reached significance. We failed to observe similarly significant differences in consumption of aversive tastes (*F < 1*; **Figure 2A-right**), but this is likely because of a floor effect on consumption of aversive tastes—note that the overall pattern of pleasant taste consumption (i.e., metestrus>estrus=proestrus>diestrus) is mirrored by an inverse pattern for aversive tastes (metestrus<estrus=proestrus<diestrus; inset of **Figure 2A-right**). This observation will be expanded upon and addressed below.

While not all rats contributed data for all estrous phases (because of occasional inter-rater disagreement on vaginal cytology, and the fact that metestrus, which persists for less than the 24 hours separating sessions, was occasionally missed), we were able to perform within-subject tests using the subset that did so. The results of these tests qualitatively replicated (data not shown) those of the completely randomized ANOVAs—the preference for pleasant tastes is higher during metestrus than in phases marked by high estradiol levels (diestrus and proestrus).

Further confirmation came from analysis of the length of licking “bouts”—rhythmic trains of licks, the lengths of which serve as a particularly sensitive measure of taste palatability (Davis & Smith, 1992; Dwyer, 2012; Lin et al., 2014; Monk et al., 2014). Bouts are easily identified because they are separated by pauses that are ≥2 times the normal inter-lick interval (ILI; note that > 95% of ILIs are within 40 ms of the 120 ms average ILI; **Figure 2B**). As these parameters of licking are consistent across estrous phase (*Kolmogorov–Smirnov D*(58190) = 0.0025, *p* = 0.72 for differences between ILI distributions), bout length analyses for different phases can be validly compared. Doing so using ANOVA revealed a significant main effect of estrous phase in pleasant taste consumption (*F*(3,146)=3.29, *p* < 0.05), an effect that again appeared to reflect a specific difference between diestrus and metestrus (*p* < 0.05; **Figure 2C-left**). As was true of mean lick counts, the overall pattern of lick bout length across estrous phase for pleasant tastes was the inverse of the trend found for aversive tastants (**Figure 2C-right inset**), but once more a floor effect (likely) kept differences in bout lengths for aversive tastes from reaching significance (*F*(3,69)=1.43, *p* > 0.05).

### Between-phase differences in preferences reflect metestrus-specific enhancement of the normal palatability range

These above analyses suggest that a small but reliable enhancement of preference for pleasant tastes emerges during metestrus, and reveal a trend toward an associated inverse effect for aversive tastes. Although the evidence for the latter effect is less compelling, we questioned the simple interpretation—i.e., that estrous cycling affects only perception of pleasant tastes—for three reasons: 1) because the inverse pattern for pleasant and aversive tastes was striking; 2) because the floor effect on consumption of aversive tastants likely blunted the “observability” of any shifts across the cycle; and 3) because the difference between the average number of licks for pleasant and aversive tastes is so large that it renders it difficult (and inappropriate) to attempt direct comparisons using a 2-way ANOVA for tastant valence (pleasant v aversive) and estrous phase.

To address these issues, we normalized the data for each taste to the average (across the entire estrous cycle) licking of that specific taste (see Methods). Thus normalized for general and individual variability, between-phase differences in licking were easily comparable, a fact that also made it possible to analyze data from both valences together, phase-by-phase, and thereby to directly test the hypothesis motivated by a comparison of the left and right panels of **Figures 2A** and **2C**—namely, that preferences magnify and diminish holistically (i.e., that enhanced preferences for pleasant tastes are accompanied by enhanced aversion of aversive tastes) with estrous phase. **Figure 3A**, a scatterplot of normalized licking to strongly valenced tastes (we focused on sucrose and quinine, because consumption of tastes that were neither strongly pleasant or strongly aversive would not be expected to change with preference magnification) supports this hypothesis, in that it reveals a significantly negative correlation between within-session normalized palatability of pleasant and aversive tastes *(r* = -0.43, *p* < 0.05). This correlation was in fact negative for 6 of 8 rats that licked appreciably to both tastes, and results were essentially the same when other highly valenced tastes (saccharin, NaCl; see Methods) were included in the analysis. Thus, we conclude that across phases of the estrous cycle, increased licking to pleasant tastes was reliably mirrored by decreased licking to aversive tastes. That is, the impact of hormonal changes on palatability is wholistic—it magnifies and shrinks the range of preferences, rather than simply changing preferences for particular tastes.

**Figure 3.**
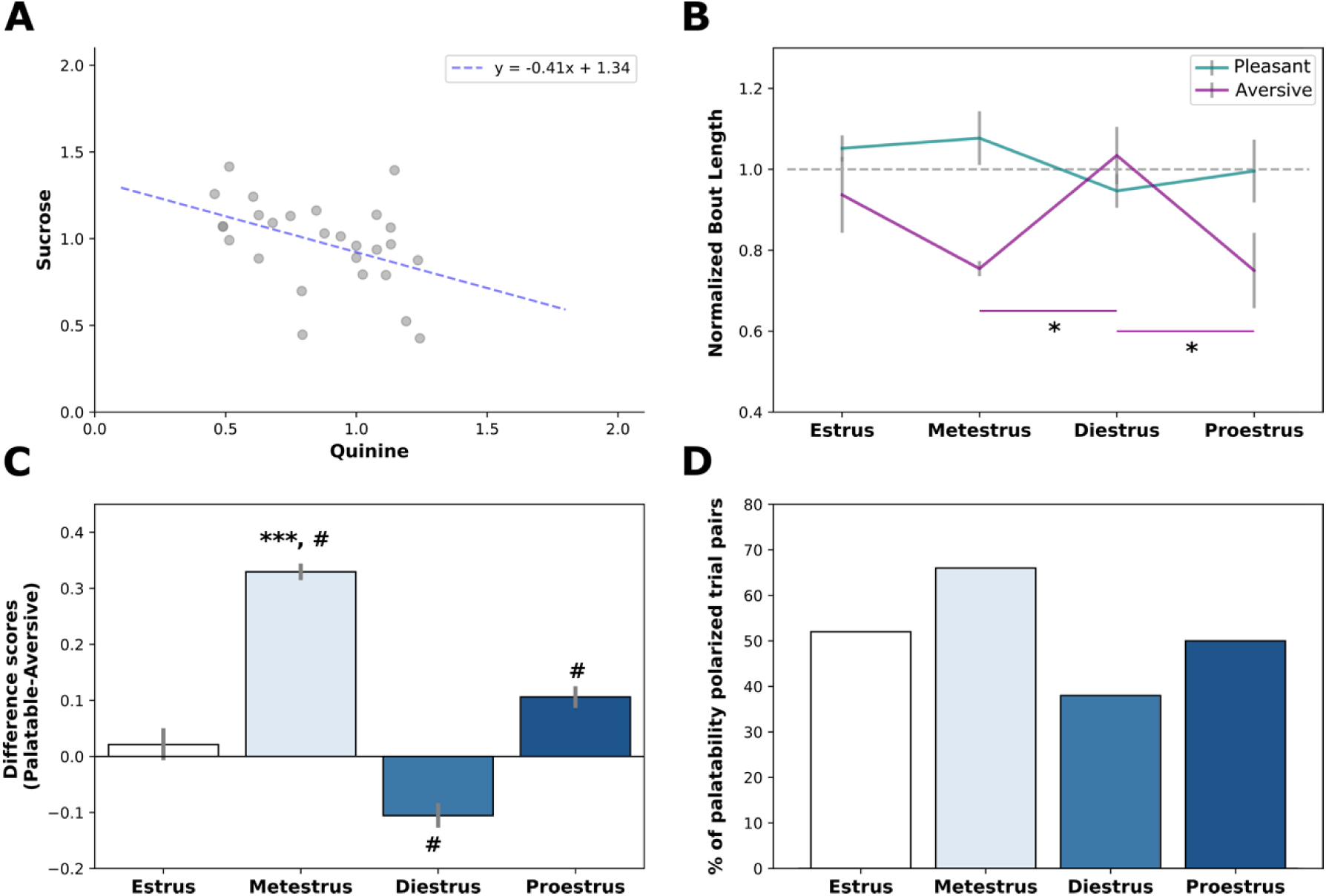
Palatability is magnified during metestrus. **A.** A scatter plot of normalized bout lengths to highly aversive (quinine) and highly pleasant (sucrose) tastes. The blue dashed line is the regression line. The significant negative correlation (p < 0.05) indicates that increases in the palatability of preferred tastes co-occur with decreases in the palatability of aversive tastes. **B**. Quantified as normalized bout lengths, preferences for pleasant tastes are elevated during metestrus, and preferences for aversive tastes are simultaneously reduced during metestrus (as well as during proestrus) relative to diestrus (* = *p* < 0.05). **C**. Difference scores calculated between pleasant and aversive tastes (see text for details) confirmed the amplification of taste palatability (y-axis) during metestrus. **D**. Percentage of pleasant-aversive trial pairs displaying palatability polarization performance across each estrous phase. *** *p* < 0.001 higher than other groups, # *p* < 0.05 different from 0. Error bars represent SEMs.

We next performed a between-phase analysis, which confirmed that the rats’ preference for pleasant tastes is maximally elevated, and their preference for aversive tastes is at its nadir, during metestrus (an effect visible because the normalization of aversive tastes was performed independently of the normalization of pleasant tastes; **Figure 3B**). A 2-way ANOVA of these data revealed a significant interaction between phase and valence (*F*(3, 52) = 3.85, *p* < 0.05) and no main effects of phase (*F*(1, 52) = 1.29, *p* > 0.05) or valence (*F*(1, 52) = 3.73, *p* = 0.06), demonstrating once again that the estrous phase-dependent effect similarly involved pleasant and aversive tastes. The perceived aversiveness of aversive tastes was significantly enhanced in metestrus (and to a smaller degree in proestrus), compared to diestrus (*ps* < 0.05); the enhancement of preference for pleasant tastes in that phase mirrored this pattern. As hypothesized, the metestrus effect appears to be a general magnification of the normal preference structure.

To further test this conclusion, we generated an unbiased estimate of the phase specificity of palatability using lick bout data, randomly sampling 50 pairs of trials between the two valences of tastes in each cycle phase and then calculating the mean differences between the pairs. Repeated 100 times, this bootstrapping process offered a direct estimate of relative magnitudes of the spread of palatability within each estrous phase. As shown in **Figure 3C**, this analysis revealed a significant (*F*(3, 296)=697.47, *p* < 0.001) magnification of the perceived difference between pleasant and aversive tastants during metestrus (*p* < 0.05): other phases either show milder (proestrus, *p* < 0.05) or no magnification (estrus), or mild diminution during diestrus (*p* < 0.05; note that more data came from diestrus than from other portions of the cycle)—in diestrus pleasant tastes remained delicious, and vice versa, but not to the same extreme as in metestrus. We performed one additional convergent test of our hypothesis, in a manner specifically designed to make the data amenable to distributional analyses (such as *X^2^*). We selected pleasant-aversive trial pairs from each session and compared the differences between the lick bout lengths in those trial pairs to the average difference for that rat across all sessions. The results of this analysis are shown in **Figure 3D**, plotted in terms of the percentage of trials in which the polarization of the two palatability extremes was larger than the average preference difference. The distribution is significantly non-uniform (*X^2^*(3) = 7.91, *p* < 0.05), with the percentage topping 50% during metestrus only. We once again conclude that the natural magnitude of the range of palatability is enhanced while female rats are in metestrus.

### Inhibition of hypothalamic estradiol binding reduces the metestrus “magnification” of palatability, but does not flatten responses across the cycle

Estradiol is the reproductive hormone most frequently implicated in feeding and related processes (Eckel, 2004; Kensicki et al., 2002; Rivera & Stincic, 2018; Wade & Gray, 1979), and thus is a reasonable candidate to explain the metestrus magnification of the palatability described above. We specifically hypothesized that this effect might reflect estradiol-driven activity in lateral hypothalamus (LH), which is: 1) involved in feeding (Anand & Brobeck, 1951a, 1951b; Margules & Olds, 1962); 2) connected to other systems involved in palatability (Berthoud & Münzberg, 2011; Fadel & Deutch, 2002; Fu et al., 2019; Petrovich et al., 2005; Tyree & de Lecea, 2017); 3) rife with neurons that produce palatability-related taste responses (Li et al., 2013); and 4) rich with estrogen receptors (Frank et al., 2014; Laflamme et al., 1998; Rivera & Eckel, 2010).

To test this hypothesis, we prepared cycling rats with bilateral guide cannulas placed just above LH (to be precise, cannulae were placed 0.1mm lateral to the center of LH, to ensure diffusion of infusate into LH while minimizing the likelihood of injection damage to the tissue itself). We then ran these rats through the same experimental protocol described above (daily preference testing followed by vaginal smears), but 20 minutes prior to each experimental session we infused either ICI_182,780_ (an estradiol α and β receptor antagonist) or DMSO (the vehicle control) into the region of LH. Using this procedure, we were able to evaluate the impact of LH estradiol inhibition on the above-described cycle-related mediation of taste palatability. Each rat was tested twice— once with each infusate, such that comparisons could be made within-rat; the order of drug was counterbalanced. Our prediction was that ICI infusions would eliminate the palatability magnification observed in metestrus, and would more generally “flatten” the cycling of palatability. **Figure 4A**, a representative coronal section stained for Nissl substance and imaged using brightfield microscopy, shows bilateral cannula tracks leading to the vicinity of LH; **Figure 4B** is the coronal schematic containing LH, overlain with dots marking the placements of infusion cannula tips for all 5 rats (one rat for which one infusion failed to target LH was removed prior to examination of licking data). Because an initial 3-way ANOVA (drug X order X estrous group) performed on licking failed to reveal significant drug order effects on preferences for pleasant or aversive tastants, we combined data across order of infusion for the analysis of preference patterning (measured in terms of normalized bout length).

**Figure 4.**
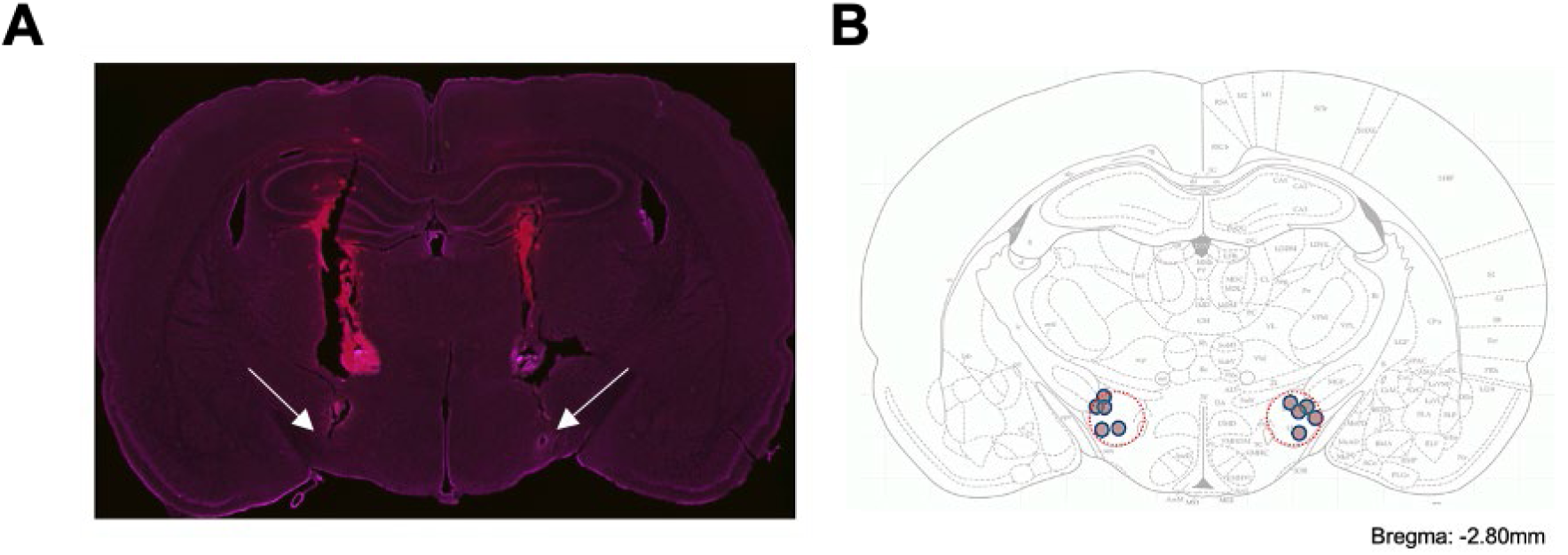
Lateral Hypothalamus (LH) cannula placement. **A**. Representative coronal section from single animal showing the placement of guide cannula. The image of the coronal section was taken at low magnification under bright-field microscopy and show Nissl Stain (magenta), placement of cannula (red), and extension of infusion cannula (2mm below guide cannula tips) in the LH. **B**. Coronal illustrations of the rat lateral hypothalamic area are depicted with the area in which bilateral cannula tips were identified in pink (each dot represents the placement of a single infusion cannula for each animal in the dataset). Correct placements were spaced from Bregma - 2.8mm according to the Paxinos rat brain atlas (Paxinos and Watson, 2006).

We first analyzed the control (DMSO-infusion) data, using the same pair of analyses brought to bear on data from non-surgical rats (i.e., Figures 3C and 3D). Both of these analyses replicated the finding of a metestrus magnification of preference extremes, with the only difference from the non-surgical results being a relatively low palatability magnification in a different (estrus) phase. While we have no conclusive explanation for this single difference (but see Discussion), it was of little statistical significance: the bootstrapping test (**Figure 5A1**; compare to **Figure 3C**) shows that normal preferences were magnified during metestrus compared to all other phases (*F*(3, 396) = 1725.86, *p* < 0.001), and the matched trials test (**Figure 5A2**; compare to **Figure 3D**) shows that the pattern of palatability magnification across the four estrous phases did not differ significantly between non-surgical and DMSO datasets (*X^2^*(3) = 6.83, *p* > 0.05). The control data adequately replicated the phenomenon.

**Figure 5.**
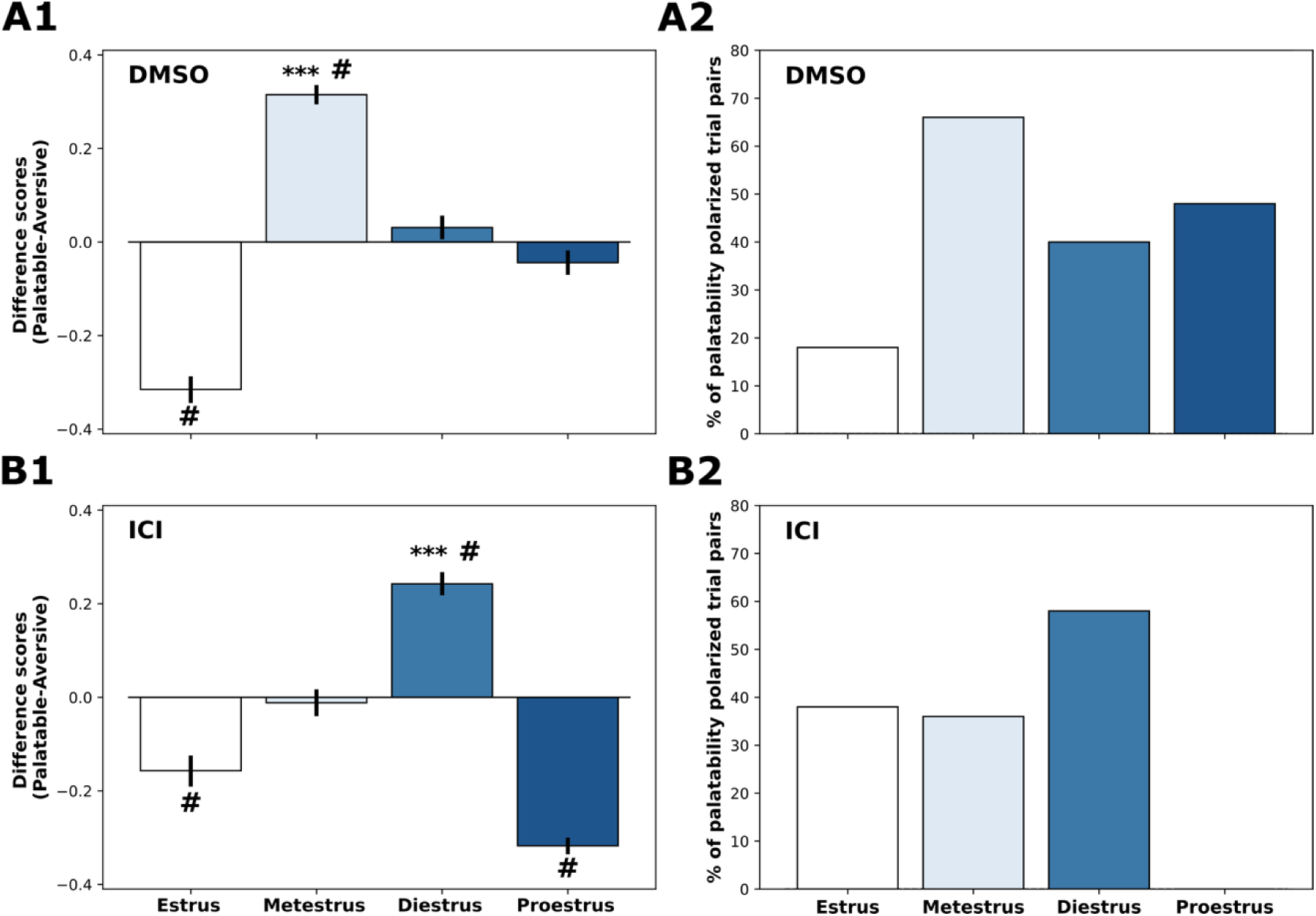
Blocking LH estrogen receptors shifts the timing of palatability preference magnification. **A.** Taste palatability polarization in metestrus characterized by preference difference scores (**A1**) and pleasant-aversive trial pairs (**A2**) during DMSO-infused behavioral sessions. **B.** Taste palatability polarization in metestrus characterized by preference difference scores (**B1**) and pleasant-aversive trial pairs (**B2**) during ICI-infused behavioral sessions. These ancillary analyses indicate that similar to the non-surgical rats (Figure 3), DMSO-treated rats exhibited the highest polarization during metestrus. Conversely, the highest polarization seemed to occur during diestrus for rats pre-treated with ICI. *** *p* < 0.001, indicating a significant magnification of taste preferences in metestrus than other phases; # *p* < 0.05 different from 0. Error bars represent SEM.

Inhibition of LH estrogen receptors, meanwhile, had a substantial impact on this taste preference pattern, although that impact was not as simple as we had hypothesized. The bootstrapping test (**Figure 5B1**) revealed (as predicted) that the infusion of ICI into LH completely eliminated the metestrus magnification of the preference differences between pleasant and aversive tastants observed in control sessions (*p* < 0.05), but further revealed that ICI infusion magnifies the normal preference patterning during diestrus (*F*(3, 396)=697.47, *p* < 0.001). It appears that LH estradiol inhibition delays, rather than eliminating, the cycle-related modulation of taste preference.

The matched pairs test (**Figure 5B2**) confirmed the impact of LH estrogen receptor blockade, showing that the across-phase pattern of palatability magnification for ICI-treated rats was significantly different from those of both the non-surgical group (*X^2^*(3) = 25.71, *p* < 0.05) and the DMSO group (*X^2^*(3) = 31.55, *p* < 0.05). Note that these *X^2^* values are both more than 3 times the size of the *X^2^* comparing the two non-ICI groups, despite several differences between the rats from which Figures 3 and 5A were generated (surgery, head tethering, fluid infusion, etc.), driving home the conclusion that the metestrus phenomenon is robust and dependent on hypothalamic action of estradiol. While the fact that ICI does not flatten the pattern must wait to be explained in future work (see Discussion), the activity of estradiol in LH clearly plays a role in shaping taste preferences and palatability.

## Discussion

Gonadal hormones (e.g. androgens and estrogens) have long been regarded as primary factors driving sex differences in consummatory behavior (Clarke & Ossenkopp, 1998; McGivern et al., 1996; Mikhail et al., 2021; Zucker, 1969); for a review see ref. 1). Fluctuations of reproductive hormones, and particularly of estradiol, appear to underlie consumption variations across estrous phases. To more definitively test whether these between-phase consumption differences might reflect differences in palatability processing, we evaluated preferences for a battery of tastes in (almost) real time, acquiring measurements at each phase of each rat’s estrous cycle. Our analysis of lick bouts revealed a general magnification of normal taste palatability patterns— preferences for pleasant tastes were at their highest, and aversions to unpleasant tastes were at their lowest—during metestrus. These results represent novel evidence that modulations in food consumption across the estrous cycle are driven by the impact of sex hormones on taste palatability, and show the phenomenon to be a generalized impact on the magnitude of normal preferences, rather than a specific manipulation of the palatability of particular tastes.

The pattern observed while examining during natural fluctuations of sex hormone across estrous phases—magnification of preference patterns in the low-estradiol metestrus phase compared to other phases—expands upon research showing that ovariectomy increases both intake and palatability of tastants in a manner that is counteracted by estrogen replacement treatment (e.g., Curtis et al., 2004; Pereira Jr et al., 2019; Zucker, 1969). It is difficult to reconcile with Atchley et al. (2005), however, who, using the same task employed here, showed a reduced licking of sucrose in estrus compared to diestrus (metestrus and proestrus licking were not reported). We saw no significant difference between licking in these two phases, a difference that could conceivably reflect concentration choice—the biggest estrus-diestrus difference observed by Atchley et al was obtained with a much lower concentration of sucrose than that used here. Alternatively, however, the observed diestrus enhancement observed by Atchley et al. might reflect the “tail end” of the metestrus enhancement that we are reporting.

It is also possible that our delivery of tastes ranging from highly pleasant to unpleasant in single sessions may have influenced the rats’ behavior, thereby causing a difference between ours and this earlier work. As noted earlier, taste palatability is intrinsically a comparative measure—an animal’s preference evaluation is dependent on the nature of the other tastes available (Ballintyn et al., 2023; Flaherty et al., 1995; Flaherty & Rowan, 1986; see also Figure 3A herein). It is, therefore, not entirely surprising to find differences in the consumption of a taste when it is delivered alone and when it is delivered among tastants of different palatabilities; the few studies (Clarke & Ossenkopp, 1998; Parker et al., 2002) that have examined cycle-dependent responses to multiple tastes have rendered results that are in broad accord with our own.

We are not the first study to identify a palatability-related phenomenon centered on metestrus. Both Fantino and Brinnel (1986) and Ter Haar (1972) also reported intake enhancements localized to that cycle phase. The fact that the largest cycle-related modulation of behavior might occur during a phase with limited estradiol circulation suggests that a full accounting of the mechanism underlying sex hormone-dependent modulation of food consumption will be temporally and hormonally complex, but is consistent with other work suggesting that the impact of a proestrus estradiol surge is often delayed by 24-48 hours (Gray & Greenwood, 1982; Santollo et al., 2007)—a theory proposed and explored by Asarian & Geary (2013). Regardless, our results jibe well with the off-cited hypothesis that food intake and taste palatability are regulated by the binding of estradiol to receptors in lateral hypothalamus (LH; see Li et al., 2013; Santollo & Daniels, 2015a, 2015b), in that intra-LH infusions of an estradiol receptor antagonist (ICI) curtailed the metestrus polarization of palatability.

Unpredicted, however, was the finding that inhibition of LH estrogen receptor activity appeared to delay, rather than completely eliminate, cycle-related fluctuations of palatability. While we have no definitive explanation for this result, it may reflect the fact that estradiol is only one element of a complex driver of taste preference structure. Given the complexity of the hormonal environment, palatability is likely the product of an interplay between multiple hormones known to fluctuate across the estrous cycle (a list that prominently includes, but is not limited to, luteinizing hormone and progesterone). If hormones naturally interact such that changes in one affect the levels of others, then a single-hormone treatment may not restore normal function in ovariectomized rats (in which levels of other hormones are also greatly suppressed). This prediction aligns with findings demonstrating that estrogen replacement treatment fails to completely mitigate the impact of ovariectomy on feeding behavior (e.g., Lampert et al., 2013). Thus, our findings highlight the value of studies looking at natural fluctuations, which enhance the interpretability of ovariectomy experiments, and also the need for more complex theories regarding the regulation of food intake by sex hormones.

It is also important to address the fact that intracranial infusions of DMSO had a non-zero impact on taste preference (compare Figure 3C to Figure 5A1), reducing in the difference between pleasant and aversive preferences (indicated by negative difference scores) during the estrus phase of the cycle. At present we are unable to say whether this estrus-phase reduction, which was also observed in rats treated with ICI, is related to some impact of DMSO itself or is an artifact of liquid infusions into LH, which potentially disrupted LH activity enough to make the rats less sensitive to taste palatability when estradiol circulation is at its lowest. Future experiments involving chemogenetic inhibition rather than intracranial infusions will test these possibilities. But regardless, this effect did not obscure the magnification of palatability during metestrus, which shifted to diestrus following the blocking of LH estrogen receptors.

In summary, our results support the hypothesis that a modulation of taste palatability likely mediates the impact of estrous phase on food consumption. Thus, palatability depends not only on taste valence, but also hormonal environment, in a manner that can only be truly appreciated in a naturalistic, within-subject experimental setting. The complex interaction between taste perception and sex hormones reflected in our data sheds light on the mechanisms underlying food craving or altered eating patterns observed following the halting of the menstrual cycle in women taking oral contraceptives (Bancroft & Rennie, 1993; Freeman et al., 2001; Marr et al., 2011), and following alterations of the menstrual cycle caused by menopause and abnormal conditions (e.g., eating disorders, excessive exercise; Coppi et al., 2021) in humans.

## Methods

### Animals

Subjects were 15 Female Long Evans rats purchased from Charles River Laboratories and housed individually from the day of their arrival until the conclusion of the experiment. Rats were kept on a 12/12 light/dark cycle, given a few days to acclimate to their new environment with free access to food and water before starting experiments, and placed on restricted water access (15mL every 24hr) to motivate participation during experiments. Their weights, which ranged from 250 to 320g on the first day of experimentation, were monitored throughout the duration of experimentation to evaluate basic health; each rat was handled each day to ensure that they were familiar with the researcher. All animal protocols were overseen and approved by the Institutional Animal Care and Use Committee (IACUC).

### Equipment

The biggest challenge in this research is the need to assess, under naturalistic conditions, the palatability of tastants within a single session—that is, within a single, one-day-long phase of the estrous cycle. The brief access task (BAT) was chosen specifically because it allows us to quantify preferences for/against multiple tastants within a single test session (lasting ∼ 30 to 60 minutes) by directly measuring licking (e.g., Caras et al., 2008; Sclafani, 2006) at spouts available for only 5-10 sec at a time.

The BAT was performed on the Davis rig MS-160 lickometer apparatus (Med Associates Inc., St. Albans, Vermont), which is comprised of a clear plastic cage and a movable platform holding multiple drinking bottles (each containing one taste stimulus). Rats can access these bottles one at a time, via an oval hole in the center of the front panel of the cage, that is available only briefly and blocked by a shutter during inter-trial intervals. Prior to each (10 sec) drinking trial, the lick spout of one bottle is moved in front of the hole, such that the opening of the shutter allows the rat to lick from that taste stimulus. Individual licks completed a (low-current) circuit, such that the Windows PC controlling the process could register each lick time.

### Taste Stimuli

Tastants used in this experiment included water (neutral), 0.3M sucrose (caloric and sweet), 0.005M saccharine (non-caloric and sweet), 0.1M sodium chloride (NaCl, salty), 0.1M citric acid (sour), and 0.001M quinine hydrochloride (QHCl, bitter), all purchased from Thermo Fisher Scientific with ACS Grade (save saccharin, which was FCC/USP Grade). Tastes and concentrations were chosen such that each was either clearly pleasant (sucrose, NaCl, saccharine) or aversive (QHCl and citric acid; e.g., Monk et al., 2014; Sadacca et al., 2012). Each tastant was dissolved and mixed in deionized water, generated in a Millipore Direct-Q® (Burlington, MA). Tastants were made fresh, the morning of each test day.

### Vaginal Cytology and Microscopy

To ensure that estrous phase was known at the time that consummatory behavior was assayed, vaginal lavages took place immediately following each (habituation and) testing session, which themselves happened at the same time every day (see below). By collecting vaginal smears after testing sessions, we avoided causing the rats undo stress prior to assaying behavior; research suggests that cytology is most validly indicative of the estrous phase during the day leading up to sample collection (Eckel & Geary, 1999).

Samples were acquired by gentling grasping and lifting the animal’s tail, exposing the ventrum and allowing the experimenter access to the entrance of the vaginal canal with a transfer pipette filled with room-temperature 0.1% PBS. The PBS was gently expressed, recollected, and placed in well plates. Two such samples were collected per rat, per session, and left to set for 20min, after which excess PBS was removed. Vaginal cells in the wells were then stained using 250µL of a 0.001M DAPI solution left on for 5min, after which 250µL of fresh PBS was added, and cells were allowed to settle for another 20 min before imaging.

Images were obtained using a brightfield overlain with a blue fluorescence filter; this allowed visualization of the number of nucleated cells in the sample. Images were captured using both 10X and 40X magnification, in order to visualize the specific cell types (40X) as well as their distribution (10X) in the sample. Estrous cycle phase was estimated by two trained coders, based on previously determined ratios of three types of cells: leukocytes, cornified, and nucleated (Cora et al., 2015). Coders were blind to both the behavioral data and the judgment of the other coder.

### Experimental Design

**Table 1** summarizes the two experiments performed for this paper. Experiments began between 9:00am and 1:00pm each day and started with five days of habituation to the BAT rig. On Habituation Day 1, rats were put in the rig with the shutter closed for 30mins, to allow them to grow accustomed to the testing space. On Day 2, the shutter was open for the entire 30mins, allowing *ad lib* access to a spout with water. On Day 3, access to water (i.e., shutter open time) was extended to 1hr. On Days 4 and 5, the shutter was only open for 10-sec trials, with the timer starting with the first lick; if the rat failed to lick for 60 sec, the shutter dropped. Shutter closing was followed by an inter-trial interval (ITI) of 30 seconds.

**Table 1.** Experimental Design.

|  |  | Days: 1-5 | Days: 6-10 | Days: 11-15 |
| --- | --- | --- | --- | --- |
| Estrous phase and behavior |  | Habituation + Lick Training | Tastes |  |
| LH estradiol and behavior | Group A | Habituation + Lick Training | DMSO + Tastes | ICI + Tastes |
|  | Group B |  | ICI + Tastes | DMSO + Tastes |

Following habituation, preference/palatability testing commenced—5 testing sessions (1/day) identical to Days 4 & 5 of habituation (e.g., lick spout available for 10 sec following first lick and 30 sec ITI), but with the rat sampling the 6-taste battery rather than just water. Each session consisted of 8 blocks of 6 trials; all 6 tastants were delivered (in randomized order) in each block. Lick and taste delivery data were saved to PC.

Once a session was complete, vaginal smears were collected, after which the rat was returned to her home cage. The rig and bottles were cleaned after each session.

### Intracranial cannula implantation and infusions of estrogen receptor blocker

Following 5 days of handling, a new set of rats underwent surgical implantation of custom-designed guide cannulas (Plastics One, Roanoke, VA) into LH. Each rat was initially anesthetized with isoflurane followed by an intraperitoneal injection of a ketamine/xylazine mixture (100/5 mg/kg); periodic intraperitoneal injections of the ketamine/xylazine cocktail (25% of the induction dose) were used to maintain stable anesthesia throughout the surgery. The animal’s head was shaved and secured in a stereotaxic frame using atraumatic ear bars, after which the skull was exposed and leveled. Support screws were attached to the skull, and bilateral craniotomies were drilled 2.8mm posterior to bregma and 2.3mm lateral to the midline; the lateral placement was chosen to avoid damaging LH by inserting infusion cannulae directly into the region; such damage may cause unwanted alterations in the consumption and welfare of rats (Morgane, 1961)(62). Cell dye (Vybrant™ Dil Cell Labeling Solution, Thermo Fisher Scientific) was applied to the tips of the guide cannulae (to aid with histological identification of tip location), which were then affixed with dummy infusion cannulae, and this assembly was slowly lowered to a depth of 6.1mm (ultimately, the infusion cannulae extended 2mm past the tip of the guide cannulae, resulting in an infusion depth of 8.1mm) below dura. Silicone was used to seal any gaps between cannulae and skull, after which dental cement fixed the entire assembly in place. A dust cap was placed over the cannula to prevent obstruction/infection, and rats were given post-operative injections of lactated ringer, penicillin, and meloxicam.

Infusions were performed after the rat was removed from their home cage prior to behavioral testing sessions. The rat was cradled in the experimenter’s lap, and infusate was delivered (total bilateral volume, 1.0 µL, delivered through cannulae threaded with 10µL Hamilton syringes) over a two-minute period. Infusate included either 80nM of ICI182,780 hydrochloride (ICI; Sigma-Aldrich), a potent α and β estrogen receptor (ER) antagonist, or Dimethylsulfoxide (DMSO; Sigma-Aldrich) vehicle. Following the infusion, the infusion cannula was left in place for another minute to allow for complete diffusion of the infusate.

After the animal was returned to its home cage for 20 minutes to ensure binding of ICI to receptors (de Arruda Camargo et al., 2000), it was moved to the BAT rig for a testing session identical to those described above. One infusate was used for 5 consecutive sessions, after which the other was used for another 5 sessions; order was counterbalanced and found to have no impact on performance (see Results). Sessions were followed by vaginal smear collection.

### Licking microstructure analysis

Licking, a highly rhythmic behavior (∼8Hz in rats), occurs in relatively discrete bouts separated by pauses of ≥2x the normal inter-lick interval (Davis, 1996; Davis & Smith, 1992); we set our inter-bout pause criterion to 0.5sec, as is common in the field (Davis, 1996; Davis & Smith, 1992; Lin et al., 2014; Spector & St. John, 1998). Bout length (i.e., the average number of licks in a bout) has been shown to be a reliable marker of preference/palatability, with longer bout lengths equating to higher palatability and smaller bout lengths equating to lower palatability (and even aversiveness; (Dwyer, 2012; Lin et al., 2014; Monk et al., 2014)).

In order to facilitate comparisons between tastants that drove vastly different bout lengths (i.e., sucrose vs quinine), we normalized each animal’s bout length using the following equation:

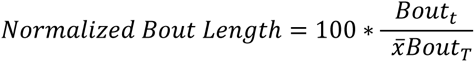

Where *Bout_t_* represents the bout length occurring during a trial of a given taste and *xBout_T_* is the mean bout length of all trials across all estrous phases of that taste. By normalizing to the mean values, we can further reveal how licking performance differs across estrous phases within tastants of each valence, thereby opening a window to investigate whether the impact of sex hormones on feeding is taste valence-specific.

### Data analysis

All statistical analyses (ANOVAs, *t*-tests, and Pearson’s correlations) were performed using custom Python scripts, with significance level (*p*) set to be < 0.05. Most were performed on tastes collapsed across palatability; correlation (r) analyses were performed only on highly valenced tastes. Significant main effects or interaction were followed by Bonferroni post-hoc tests.

### Histology

After completing the experimental protocols, rats were deeply anesthetized with a ketamine/xylazine mixture (200/10 mg/kg) and exsanguinated *via* cardial perfusions of 0.9% saline and 10% formalin. Brains were extracted and fixed in a 30%/10% sucrose/formalin solution for two days. After the brains were fixed, coronal slices (60 µm) were made, mounted onto slides, and stained with NeuroTrace™ (500/525 Green Fluorescent Nissl Stain, Thermo Fisher Scientific) to allow assessment of infusion cannula placements.

## Acknowledgements

This work has been supported by National Institute on Deafness and Other Communication Disorders Grants R01-DC006666 & DC007703 to DBK, R21-DC016706 to JYL and funding through the National Science Foundation (NSF Grant #: IBN180002 to DBK).

